# eQTM (expression quantitative trait methylation) Atlas: a comprehensive resource of over 11 million DNA methylation–gene expression associations through across 11 tissues and 4 diseases

**DOI:** 10.64898/2026.06.07.730721

**Authors:** Aditya Sriram, Soyeon Kim, Rebecca Caldino Bohn, Wei Chen, Tianhao Liu, Molin Yue, Niyati Jain, Brandon L. Pierce, Roby Joehanes, Daniel Levy, Etienne Patin, Lluis Quintana-Murci, Hyun Jung Park, Juan C. Celedón

## Abstract

**Motivation:** Epigenome-wide association studies (EWAS) have identified numerous DNA methylation (DNAm) CpG sites associated with complex traits and diseases, but interpretation of those CpG sites remains challenging because in EWAS, CpGs are mostly linked to nearby genes based only on genomic proximity. Expression quantitative trait methylation (eQTM) analyses connect DNAm CpGs with statistically associated gene expression levels. However, a comprehensive, searchable resource integrating eQTMs across diverse tissues and disease contexts has been lacking.

**Results:** We developed the eQTM Atlas, a web-based resource that manually curates more than 11 million DNAm–gene expression associations from eight cohorts, covering 11 tissue types, four broad disease contexts, 173,886 unique CpG probes and 20,231 unique genes. The Atlas supports gene- or CpG-searches by tissue or disease type and finding associated CpG or genes, visualization of cis- and trans-eQTMs through genome browser, heatmap interfaces across various tissues, and cohort-level data downloads. By integrating eQTM results with EWAS resources, the eQTM Atlas enables users to connect disease- or trait-associated CpGs to statistically associated genes rather than relying solely on proximity-based gene annotation, supporting functional interpretation of EWAS findings and generation of disease-specific regulatory hypotheses.

**Availability and implementation:** The eQTM Atlas is freely available at https://shiny.crc.pitt.edu/eqtm_browser/. The web interface is implemented in R Shiny and hosted through the University of Pittsburgh Center for Research Computing (CRC). Source code is available at https://github.com/ads303/eQTM-Atlas.

## Introduction

Large-scale epigenome-wide association study (EWAS) databases have become important resources for providing DNAm CpG sites associated with complex traits, environmental exposures, and diseases. These resources, including the EWAS Atlas and EWAS Open Platform, EWAS catalog, enable researchers to search disease- and trait-associated CpGs across 1908 published studies and provide a valuable foundation for downstream biological interpretation (Battram, et al., 2022; Li, et al., 2019; Xiong, et al., 2021). However, most EWAS databases annotate CpG sites to genes primarily based on genomic location, such as whether a CpG overlaps with or lies near a gene or promoter region. Although this proximity-based annotation is useful for interpreting DNAm CpG sites at the gene level, it does not necessarily identify the gene whose expression is statistically associated with DNAm at that CpG.

This limitation is important because the biological interpretation of EWAS findings often depends on the annotated gene to each CpG. Researchers often use these CpG-to-gene annotations as the basis for downstream analyses, including pathway enrichment, disease-mechanism interpretation, and target prioritization. If a CpG is assigned only to a nearby gene, researchers may miss a gene whose expression is statistically associated with DNAm level at that CpG but is located farther away, and they may prioritize a nearby gene that is not associated with the DNAm CpG finding. Therefore, EWAS findings require functional gene annotation that goes beyond genomic proximity.

Expression quantitative trait methylation (eQTM) analysis provides a data-driven approach for this purpose by linking DNAm levels at CpG sites to statistically associated gene expression levels. In contrast to proximity-based annotation, eQTM-based annotation can identify genes whose expression is significantly associated with DNAm CpG, including both local cis-associations and trans-associations. This makes eQTM results particularly useful for connecting EWAS CpGs to potential gene-regulatory mechanisms. However, eQTM associations are highly context dependent and vary across tissues and disease states (Battram, et al., 2022; Kim, et al., 2026; Kim, et al., 2020). Therefore, the same EWAS CpG may have different associated genes depending on tissue type or disease type.

Despite the growing number of eQTM studies, a comprehensive and searchable resource for tissue-specific and disease-specific gene annotation of EWAS CpGs has been lacking. As a result, researchers often continue to rely on nearby-gene annotations from EWAS databases, even when statistically associated genes can provide a more biologically informative interpretation. A centralized eQTM resource that integrates DNAm–gene expression associations across multiple tissues and disease contexts would help bridge this gap by enabling users to identify context-relevant genes associated with EWAS CpGs.

To address this need, we developed the first eQTM database, eQTM Atlas, a web-based resource that integrates manually curated DNAm–gene expression associations across diverse tissues and disease contexts (Figure 1). The current eQTM Atlas, as of June 2026, includes more than 11 million eQTM associations from eight cohorts, spanning 11 tissue types and four broad disease contexts, and covering 173,886 unique CpG probes and 20,231 unique genes (Table 1). The Atlas enables users to search by CpG or gene, visualize cis- and trans-eQTM associations, compare DNAm–gene expression relationships across tissues, download cohort-level summary statistics, and integrate eQTM results with EWAS resources.

**Table 1.** Summary of eQTM studies and corresponding tissue types included in the eQTM Atlas.

| Study | Disease | # of subjects | eQTM Pairs | Tissue | Reference | Cis/Trans |
| --- | --- | --- | --- | --- | --- | --- |
| **EVA-PR** | atopic asthma | 158 | 103,356 | Nasal Epithelium | (Kim, et al., 2023) | Cis and trans |
|  | non-atopic healthy control | 100 | 1,518 | Nasal Epithelium | (Kim, et al., 2023) | Cis and trans |
|  | asthma + non-asthma | 455 | 9,078,542 | Nasal Epithelium | (Kim, et al., 2020) | Cis and trans |
|  | atopic asthma + healthy controls | 258 | 2,140,608 | Nasal Epithelium | (Kim, et al., 2023) | Cis and trans |
|  | White blood cell | 434 | 157,044 | White Blood Cell | (Kim, et al., 2023) | Cis and trans |
| **FUSION** | T2D & Diabetes-Related Traits | 265 | 36,516 | Skeletal Muscle | (Taylor, et al., 2019) | Cis and trans |
| **Framingham Heart Study** | cardiometabolic disease | 2115 | 10,000 | Whole Blood | (Keshawarz, et al., 2023) | Cis |
| **Colonomics** | colon cancer | 132 | 165 | Colon | (Diez-Villanueva, et al., 2021) | Cis |
| **EvoImmunoPop** | healthy | 156 | 811 | Primary Monocyte | (Husquin, et al., 2018) | Cis |
| **HELIX** | healthy children | 832 | 32,677 | Whole Blood | (Ruiz-Arenas, et al., 2022) | Cis |
| **GTE<sub>x</sub>** | healthy | 424 | 6,318 | Breast Mammary | (Oliva, et al., 2023) | Cis |
|  | healthy | 424 | 6,650 | Transverse Colon | (Oliva, et al., 2023) | Cis |
|  | healthy | 424 | 8,686 | Lung | (Oliva, et al., 2023) | Cis |
|  | healthy | 424 | 4,319 | Skeletal Muscle | (Oliva, et al., 2023) | Cis |
|  | healthy | 424 | 9,251 | Ovary | (Oliva, et al., 2023) | Cis |
|  | healthy | 424 | 6,203 | Prostate | (Oliva, et al., 2023) | Cis |
|  | healthy | 424 | 2,439 | Testis | (Oliva, et al., 2023) | Cis |
|  | healthy | 424 | 3,916 | Whole Blood | (Oliva, et al., 2023) | Cis |

**Figure 1.**
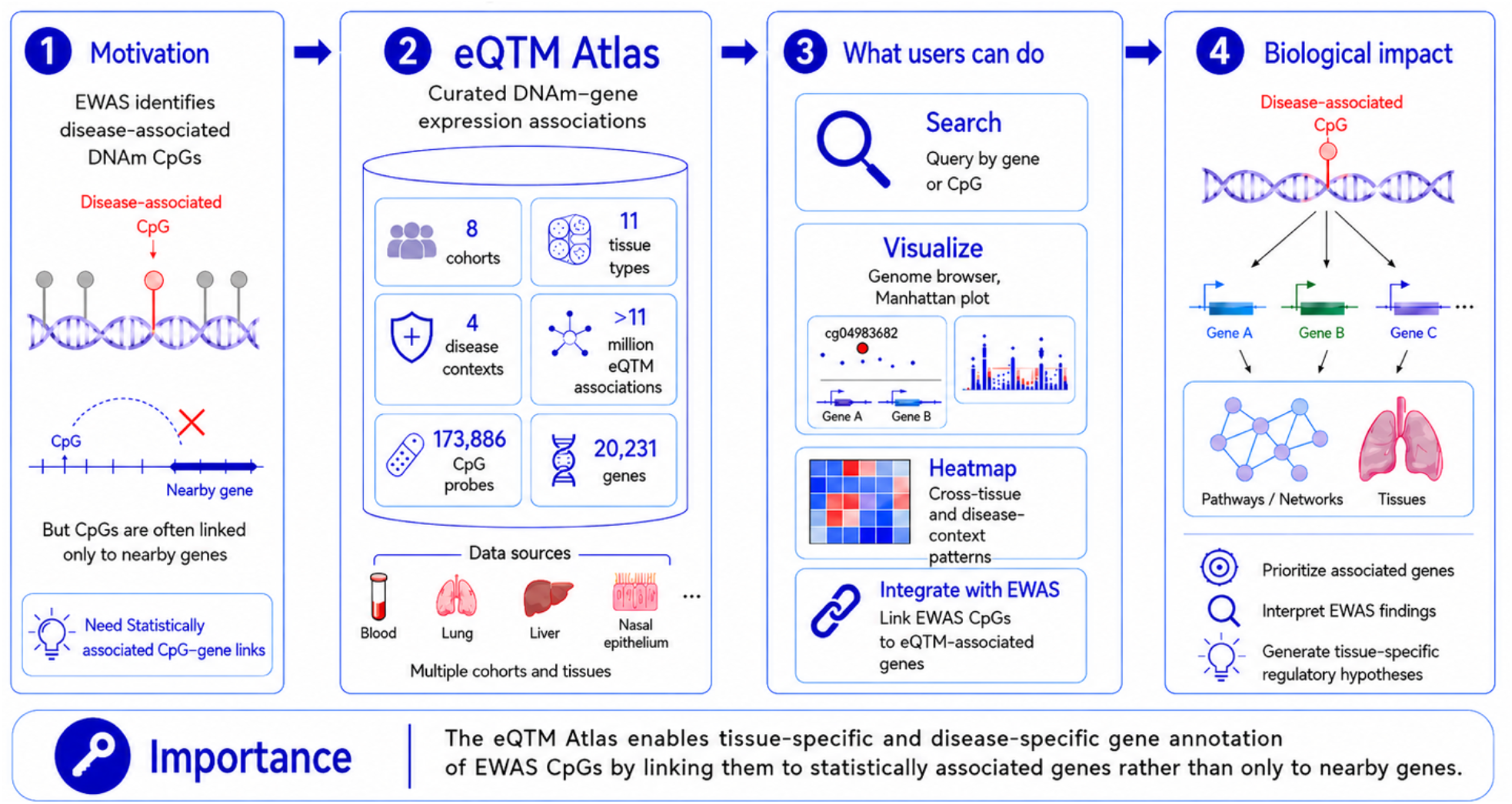
Overview of the eQTM Atlas. The eQTM Atlas is a curated resource of DNAm– gene expression associations. The Atlas integrates eQTM results from eight cohorts, 11 tissue types, and four disease contexts, including more than 11 million eQTM associations, 173,886 CpG probes, and 20,231 genes. Users can search by gene or CpG, visualize associations using genome browser and Manhattan plot views, examine cross-tissue patterns using heatmaps, and integrate EWAS findings with eQTM-associated genes to support tissue- and disease-specific gene annotation and regulatory hypothesis generation.

A key contribution of the eQTM Atlas is that it enables tissue-specific and disease-specific gene annotation of EWAS CpGs by linking them to statistically associated genes rather than nearby genes alone. By connecting EWAS-associated CpGs with eQTM-associated genes across relevant biological contexts, the eQTM Atlas could support functional interpretation of EWAS findings, prioritization of disease-relevant genes, and generation of tissue-specific regulatory hypotheses.

## Materials and methods

### Data summary of eQTM Atlas

The eQTM Atlas provides a comprehensive resource summarizing published and newly generated eQTM studies. Through manual curation and standardized formatting, the atlas currently includes eight eQTM studies, representing approximately 4,379 individuals across 11 tissue types and four disease types (Figure 1, Table 1). Collectively, the eQTM Atlas contains more than 11 million eQTM associations, corresponding to 173,886 unique CpG probes and 20,231 unique genes. We collected both cis- and trans-eQTMs (Figure 2). Cis-eQTMs were defined as associations in which the associated gene is located near the CpG site, typically within 1 Mb of the transcription start site (TSS), whereas trans-eQTMs were defined as associations in which the associated gene is located farther away, either on the same chromosome or on a different chromosome (Keshawarz, et al., 2023).

**Figure 2.**
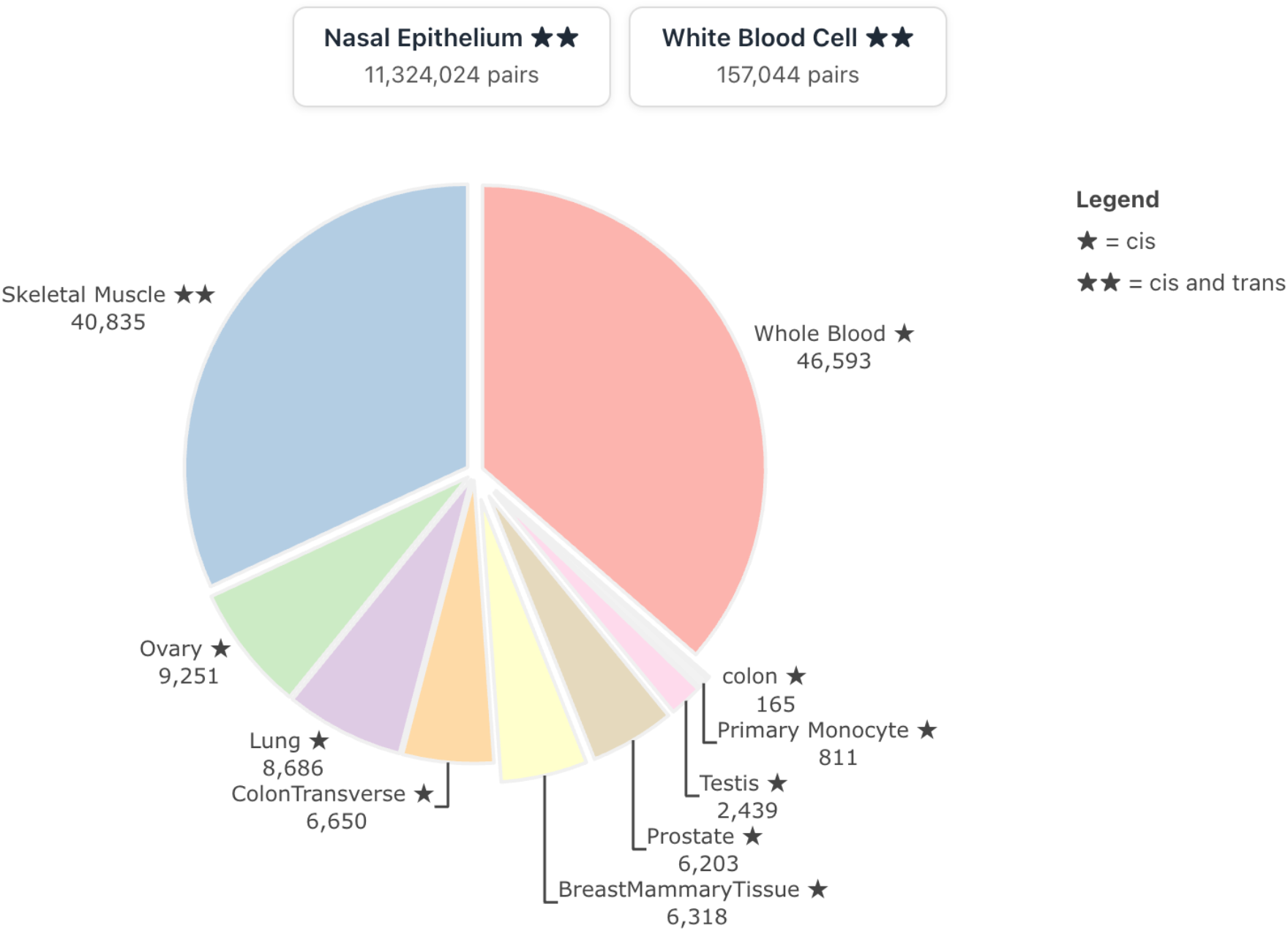
Distribution of curated eQTM pairs by tissue in the eQTM Atlas. The pie chart shows the number of curated CpG–gene eQTM pairs for each tissue. Stars indicate eQTM type: ★, cis only; ★★, both cis and trans. Nasal epithelium and white blood cells are shown separately above the pie chart because their large numbers of eQTMs would otherwise dominate the tissue distribution.

Our database includes 8 cohorts. A particularly significant initiative is the Genotype-Tissue Expression (GTEx) Consortium, which has comprehensively mapped cis-eQTM across eight different tissue types, including breast, colon, lung, skeletal muscle, ovary, prostate, testis, and whole blood(Oliva, et al., 2023). We also included other studies in various tissues, such as in colon(Diez-Villanueva, et al., 2021), monocyte(Husquin, et al., 2018), skeletal(Taylor, et al., 2019), whole blood(Keshawarz, et al., 2023), and children’s blood(Ruiz-Arenas, et al., 2022) (Table 1).

Notably, we incorporated results from our eQTM studies using the EVA-PR cohort (Epigenetic Variation and Childhood Asthma in Puerto Ricans), which includes nasal epithelium samples from children with and without asthma. We performed eQTM analyses across several subgroups: (1) atopic asthma and healthy controls (Kim, et al., 2023), (2) atopic asthma only, (3) healthy controls only, and (4) asthma and controls combined (Kim, et al., 2026; Kim, et al., 2020). A distinctive feature of these analyses is the identification of a large number of trans-eQTMs, exceeding 9 million associations, which represents a unique contribution to the eQTM Atlas. In addition, for largely overlapping individuals in these groups, we curated eQTMs from white blood cells (WBC), enabling researchers to compare eQTM profiles between nasal epithelium and WBC. This comparison is particularly relevant because both tissues are influenced by immune-related regulatory processes (Kim, et al., 2024).

### Connecting to EWAS

The eQTM Atlas also integrates one of the largest EWAS database, the EWAS Open Platform, which contains 862,697 EWAS associations across 898 phenotypes and 248 tissues/cells from 1204 publications as of May 2026 (https://ngdc.cncb.ac.cn/ewas/atlas/index). While the Open Platform annotates genes based on their physical proximity—specifically whether they reside within a gene or promoter region—the eQTM Atlas identifies genes that are significantly associated with EWAS CpGs. This distinction is critical because downstream analyses, such as pathway enrichment, frequently rely on these “nearby” gene annotations rather than on genes whose expression is actually linked to the CpGs. This reliance on proximity can lead to misleading inferred pathways that are disconnected from the underlying methylation signals.

Furthermore, the limitations of proximity-based annotation are highlighted by the fact that 16.8% of CpGs in the 850K array (151,037 out of 897,322) remain entirely unannotated because they fall outside traditional gene or promoter boundaries. By bridging these gaps, the eQTM Atlas provides researchers with a more robust, data-driven approach for downstream analysis.

### Data collection and quality control

We manually curated the complete list of significant eQTM associations, primarily by obtaining results directly from the authors of published studies, following a set of predefined inclusion criteria:

1. The studies analyzed human samples from diverse biological contexts, including normal and disease conditions.
2. We included eQTM datasets generated from genome-wide eQTM mapping. We excluded datasets from secondary analyses, such as studies that performed eQTM analysis only on EWAS CpGs.
3. To ensure adequate statistical power for eQTM analysis, only studies with a minimum of 100 samples were included.
4. Only transcriptome data generated using reliable, well-established sequencing technologies, such as Illumina NextSeq 500 and Illumina HiSeq 2000, were considered (Table S1).
5. For consistency, DNAm data were generated using either the HumanMethylation450 BeadChip array or the MethylationEPIC BeadChip (“EPIC”) array.

After thoroughly reviewing the literature and applying these criteria, we retained 18 eQTM studies in our database (Table 1). The collected data included eQTM summary statistics, including CpG, genomic coordinates, associated genes, the genomic locations of the CpGs and genes, the coefficients of the CpG–gene expression association, p-values, and adjusted p-values such as false discovery rates (FDRs).

## Results

### Web design and interface

eQTM Atlas offers a dedicated web interface that allows users to explore, discover, and contribute newly generated or curated datasets. The eQTM Atlas offers five main interactive interfaces:

i. The **‘Gene**↔**CpG Search’** feature allows users to easily search for specific genes and **CpG sites** within the database. Users can input multiple genes or CpGs and download eQTM results from various studies (Figures 3A, 4A). This tab is also accessible by searching a gene or CpG site from the ‘**Home Page’** search box.
ii. The **‘Genome Browser’** tab provides a detailed view of eQTMs in reference to their associated genomic regions, enabling users to explore the associations between **CpG sites** and gene expression in a visually interpretable way (Figures 3B, 4B).
iii. The **‘Heatmap’** tab lets users visualize the significance of eQTMs across different tissues via heatmap renderings, allowing for scalable cross-tissue comparison (Figures 3C, 4C).
iv. The **‘EWAS-eQTM Integration’** tab connects EWAS results and eQTM results, allowing users to connect CpG sites from EWAS with significantly associated genes instead of nearby genes. This tab connects eQTMs from the 8 studies we collected for the database and EWAS results from the EWAS Atlas (Li, et al., 2019) across 1,906 studies (Figure 3D).
v. The **‘Downloads’** tab allows users to download specific cohort-level summary statistics. (Figure 5A)
vi. The Upload Your Data/Site Feedback tab allows users to submit their own eQTM datasets for inclusion in the catalog, making them available for other users to download (Figure 5B).

**Figure 3.**
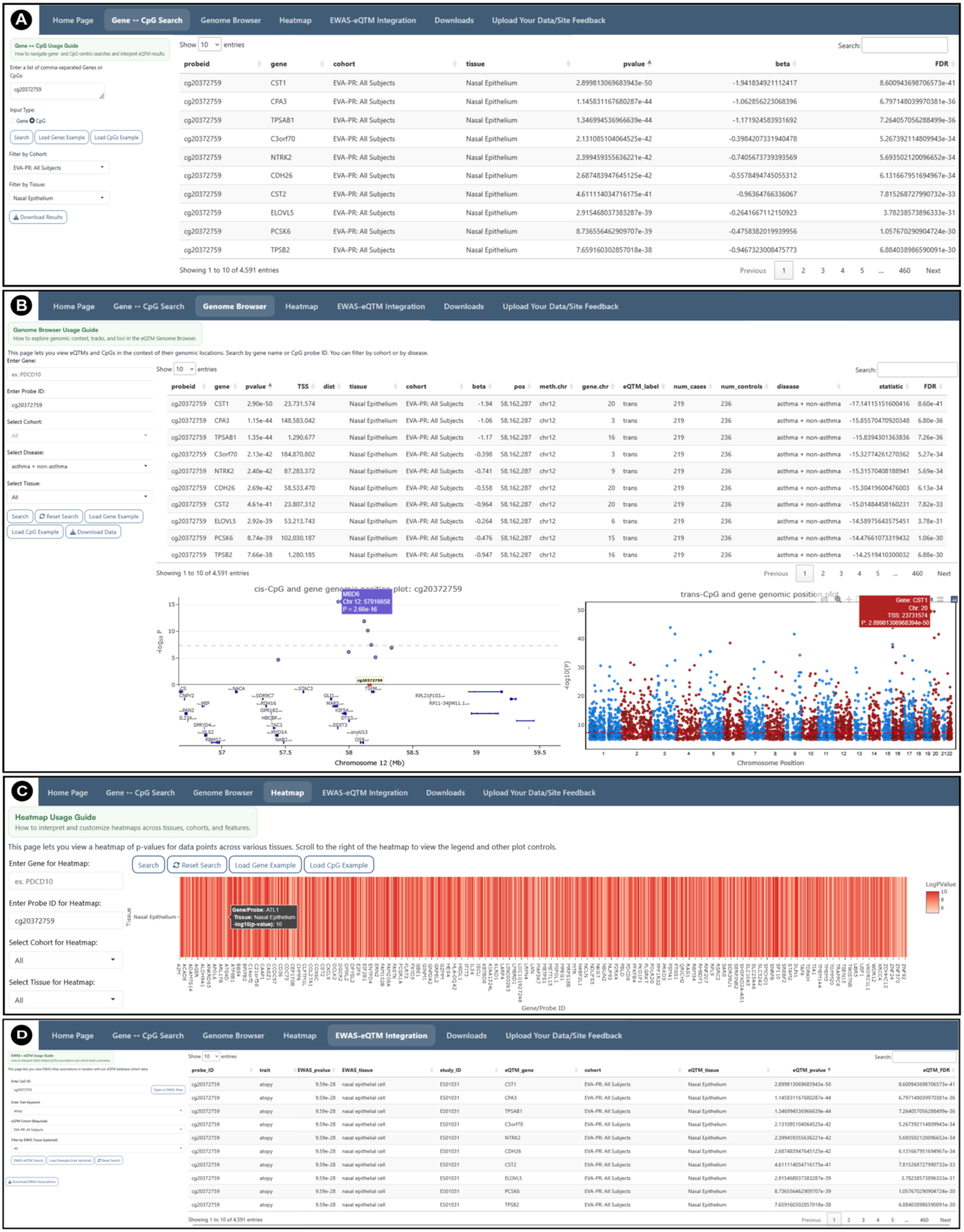
Representative database screenshots for the CpG search interface in the eQTM Atlas using **cg20372759** as an example. (A) CpG-Gene Search: A summary table of eQTM results from a multi-CpG query, filterable by cohort and tissue. (B) Genome Browser: An integrative view of a CpG’s eQTM associations, featuring a detailed results table, a genomic locus close-up showing gene annotations for cis-associations, and a Manhattan plot for trans-associations. (C) Heatmap: A p-value heatmap of a CpG’s associations with multiple genes across various tissues, providing an overview of tissue-specific regulation. (D) EWAS-eQTM Integration: A combined results table integrates associations from the EWAS Atlas and eQTM Atlas, enabling users to identify genes significantly associated with EWAS CpGs by cohort and trait.

**Figure 4.**
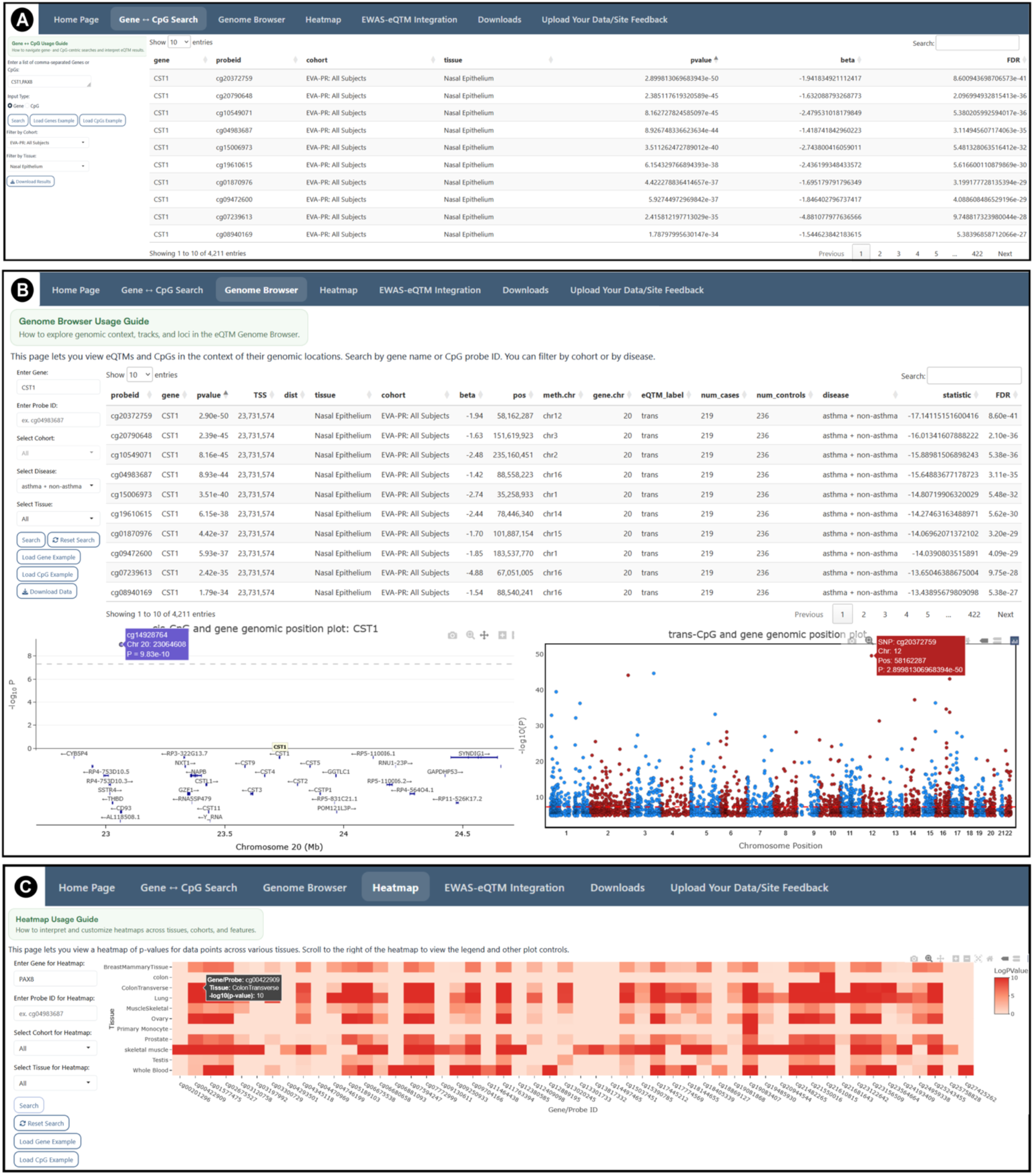
Representative screenshots for a gene search on the eQTM Atlas using CST1 and PAX8 genes as examples. (A) Gene-CpG Search: A summary table of eQTM results from a multi-gene query, filterable by cohort and tissue with interactive column sorting. (B) The Genome Browser: An integrative view of a gene’s eQTM associations, featuring a detailed results table, a close-up of the genomic locus with gene annotations for cis-associations, and a Manhattan plot for trans-associations. Results are filterable by cohort or disease. (C) The Heatmap: Displays a p-value heatmap for a gene’s associations with different CpGs across various tissues, providing an overview of tissue-specific associations.

**Figure 5.**
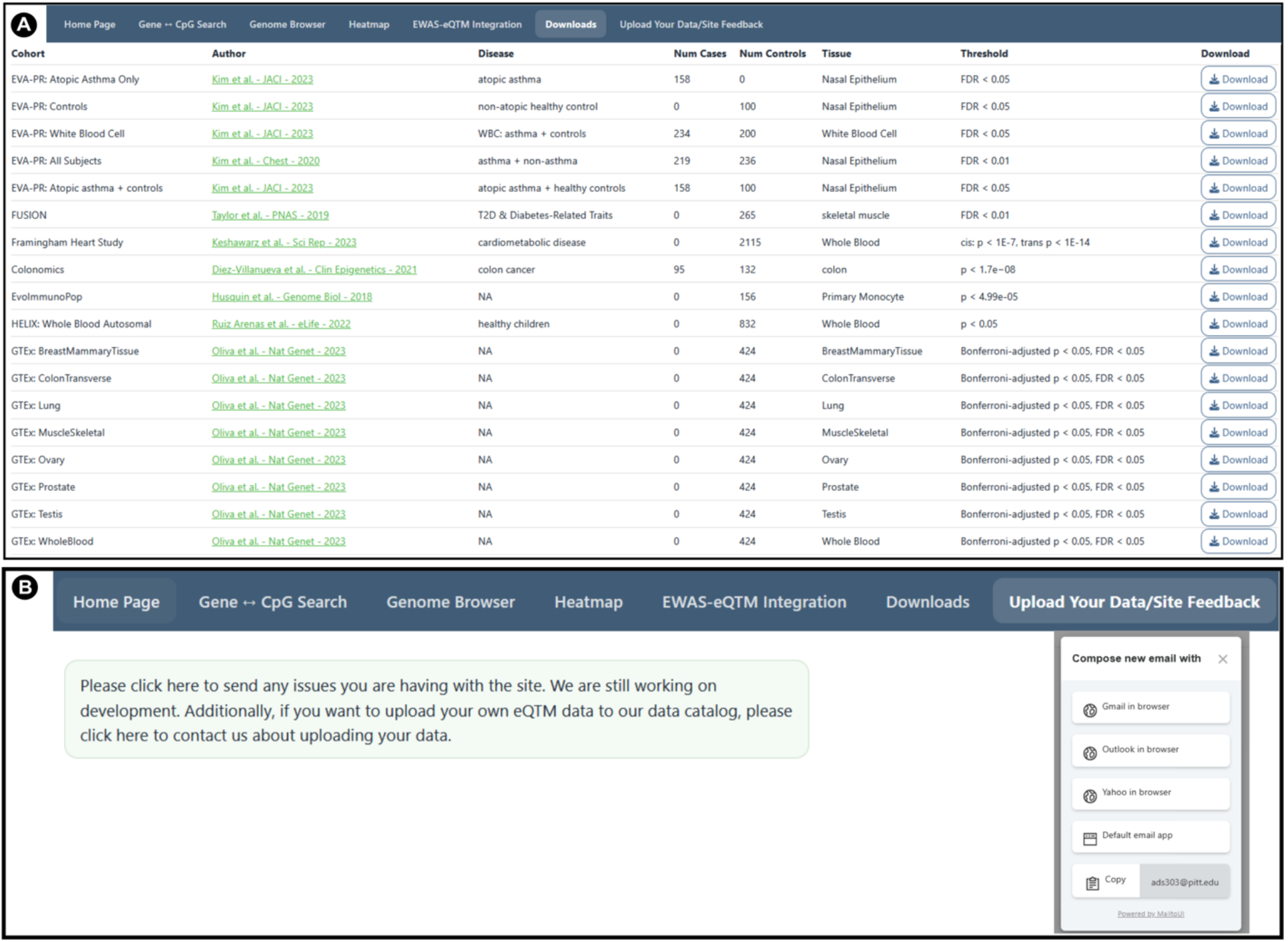
Data download and user-feedback tabs of the eQTM Atlas. (A) The Downloads page provides a cohort-level catalog of curated eQTM datasets included in the eQTM Atlas. Users can directly download eQTM results from various tissues and cohorts (B) The Upload Your Data/Site Feedback page enables users to contact the development team to report website issues or contribute additional eQTM datasets.

### CpG search example

To illustrate a typical use case of the eQTM Browser, we queried **cg20372759**, the CpG site most significantly associated with atopy the EVA-PR cohort (Forno, et al., 2019). In the **“Gene ↔ CpG Search”** tab (Figure 3A), users can enter a CpG ID, such as cg20372759, and optionally restrict the search by cohort and tissue—for example, the EVA-PR cohort and nasal epithelium.

The resulting table displays all genes whose expression is significantly associated with the queried CpG, along with the corresponding cohort, tissue, effect size (β), P value, and FDR, sorted by statistical significance. The full results table can be downloaded as an Excel file, and the associated gene list can be used for downstream analyses, including pathway or gene-set enrichment analysis. Users may also paste multiple CpG IDs to retrieve CpG-associated genes in batch.

In the **“Genome Browser”** tab (Figure 3B), users can visualize the genomic locations of genes associated with a queried CpG, together with the statistical significance of each association, shown as –log10(*P* value). Cis- and trans-associations are displayed separately. The cis plot shows the relative genomic positions of associated genes near the CpG, whereas the trans plot is presented as a Manhattan plot, indicating the chromosomes on which the associated genes are located. By hovering over individual data points in either the cis or trans plot, users can view gene-level details, including gene name, genomic location, and eQTM *P*-values. Although not shown in the figure, summary statistics for the genes associated with the queried CpG are provided at the bottom of the tab and sorted by statistical significance.

The **“Heatmap”** tab (Figure 3C) visualizes the −log10(*P*) values of eQTM associations between the queried CpG and its associated genes across all available tissues. This view enables users to assess whether CpG–gene associations are tissue-specific or broadly shared across tissues. For cg20372759, the associations appear to be specific to nasal epithelium, with significant associations observed across multiple genes. Users can hover over each heatmap cell to view the exact significance value for each CpG–gene association in a given tissue.

Finally, the “EWAS–eQTM Integration” tab (Figure 3D) links a DNAm CpG site to published EWAS results, helping users explore its potential epigenetic regulatory role in disease contexts.

In this tab, users can search for CpG–disease associations by entering a CpG ID and opening the corresponding record by clicking “Open EWAS Atlas” box (Figure S1).

To identify eQTM genes associated with an EWAS CpG, users can enter the probe ID, select a cohort, which is a required field, and optionally select one of the catalogued EWAS traits. In this example, we queried cg20372759, selected the EVA-PR cohort, and did not restrict the search to a specific trait. The resulting table displays 6 EWAS diseases/traits linked to cg20372759, together with their corresponding eQTM-associated genes. For cg20372759, the eQTM Atlas identified 2,112 associated eQTM genes. One of the eQTM genes, CST1, has been implicated in airway inflammation and mucus overproduction in asthma (Kim, et al., 2023) and most significantly differentially expressed gene in atopic asthma (Forno, et al., 2020). Therefore, examining its potential regulation through DNAm can provide insight into disease mechanisms. Notably, cg20372759 is associated with five diseases and one phenotype, all reported in nasal epithelial cells or airway epithelial cells. Therefore, eQTM results from nasal epithelium are particularly relevant for interpreting the potential regulatory mechanisms underlying these EWAS associations.

### Identifying eQTM DNAm CpGs through gene search

To illustrate gene-based searches in the eQTM browser, we used CST1 and PAX8 as representative examples. These genes were selected because they show distinct patterns of eQTM associations: CST1 exhibits numerous associations with DNAm CpGs primarily in nasal epithelium, whereas PAX8 shows eQTM associations across multiple tissue types. The PAX8 gene is a master regulator transcription factor essential for the embryonic development of the thyroid gland, kidneys, and Müllerian system (reproductive tract). It plays a critical role in tissue differentiation and maintenance and is used in pathology as a key biomarker for thyroid and renal cancers (Fernández, et al., 2015).

On the **“Gene ↔ CpG Search”** tab (Figure 4A), we entered *CST1 and PAX8* in the Gene field (leaving the CpG field empty) and restricted the query to the EVA-PR cohort and nasal epithelium. The resulting table lists all CpG sites whose methylation levels are associated with *CST1 and PAX8* expression, together with the cohort, tissue, effect size (β), P value and FDR, sorted by significance.

In the “Genome Browser” tab (Figure 4B), CST1 is visualized in its genomic context. The upper panel displays cis-eQTM CpGs located near CST1, allowing users to examine which nearby CpGs are associated with CST1 expression and the strength of each association, represented as – log10(P-value). The lower panel presents a Manhattan plot of trans-eQTMs across the 22 autosomes, excluding the X and Y sex chromosomes, highlighting the numerous distal DNAm CpGs associated with CST1 expression. Together, these complementary views enable users to quickly assess whether the epigenetic associations with a given gene are primarily local, distal, or both.

Finally, the “Heatmap” tab (Figure 4C) displays the significance of associations between DNAm levels at individual CpG sites and PAX8 expression across all available tissues. This visualization enables users to assess whether gene–CpG associations are tissue-specific, such as those enriched in nasal epithelium, or broadly shared across tissues. PAX8 is significantly associated with CpGs in all 11 tissues. This is consistent with its central role as a master regulator transcription factor, and therefore, epigenetic dysregulation of PAX8 could lead to diverse functional alterations across multiple tissues.

### Implementation, Hosting, and Technical Specifications

The eQTM Atlas is implemented with R Shiny, an open-source web application development tool using R code framework, locuszoomr, a web tool formatted as an R package to allow for plotting detailed and interactive gene locus plots (Pruim, et al., 2010), and shinyjs, a Javascript container within R Shiny that allows execution of powerful Java tools within the R interface. To allow for rapid data retrieval and quick plot rendering, the eQTM Atlas is hosted on the University of Pittsburgh Center for Research Computing high throughput computing (HTC) cluster.

Lastly, Users can also upload their own tissue-labeled eQTM data to our portal with an appropriate data transfer protocol (MTA/DTA) from their affiliated institution (Figure 5A). We perform a combination of automated and manual data quality control to ensure that files are all consistently formatted upon upload to our server. We currently only have open-source summary statistics available on our platform; however, we will also offer access-controlled data from investigators and individuals who provide their individual-level results for use.

## Conclusions

In conclusion, we developed the eQTM Atlas as a comprehensive resource for exploring associations between DNAm CpG sites and gene expression across diverse human tissues and disease contexts. By integrating more than 11 million eQTM associations from multiple cohorts and tissue types, the eQTM Atlas provides comprehensive resources for querying, visualizing, downloading, and comparing DNAm–gene expression relationships. Importantly, by linking EWAS-associated CpGs to statistically associated genes rather than relying only on nearby gene annotations, the eQTM Atlas complements existing EWAS resources such as the EWAS Open Platform. Instead of linking nearby genes, eQTM atlas enables researchers to prioritize genes whose expression is statistically associated with disease- or trait-relevant CpG sites. Through its searchable interface, genome browser, cross-tissue heatmaps, EWAS–eQTM integration, and downloadable summary statistics, the eQTM Atlas provides a practical platform for functional interpretation of EWAS findings, generation of mechanistic hypotheses, and identification of tissue-specific potential epigenetic regulatory relationships.

## Funding

This work was supported by grants HL079966, HL117191, MD011764 and HL168539 (to J.C.C.), and K01 HL153792 (to S.K.) from the U.S. National Institutes of Health (NIH). W.C. was supported by NIH grant HL150431 and NSF grant 2225775. H.J.P. was supported by the UPMC Hillman Cancer Center Biostatistics Shared Resource that is supported in part by award P30CA047904 and R01GM108618 at the NIH. H.J.P. is also supported by the Hillman Cancer Center Career Enhancement Program Award (P50 CA254865-01). This research was supported in part by the University of Pittsburgh Center for Research Computing, RRID:SCR_022735, through the resources provided. Specifically, this work used the HTC cluster, which is supported by NIH award number S10OD028483 and H2P cluster, which is supported by NSF award number OAC-2117681.

## Authors’ contributions

S.K. conceived and led the development of the eQTM Atlas, coordinated data curation and analysis. A.S. implemented the database, and A.S. and S.K. wrote the first draft of the manuscript; R.B., T.L., and M.Y participated in figure generation; H.J and W.C. provided the initial idea of building the database; H.J collected the summary statistics of various eQTM datasets and edited the first draft of the manuscript. L. J, B. P, R.J. D.L, E.P. L, Q, and J.C.C. provided summary statistics of one of the eQTM cohorts. J.C.C. edited the final draft; and all authors reviewed and approved the final version of the manuscript.

## Competing interests

The authors declare no competing interests.

## Availability of data and materials

The eQTM Atlas is available at https://shiny.crc.pitt.edu/eqtm_browser/ Code is available at https://github.com/ads303/eQTM-Atlas

---

**Table S1.** Number of cis and trans eQTM associations by cohort.

| Cohort | cis | trans |
| --- | --- | --- |
| Colonomics | 165 | 0 |
| EVA-PR: All Subjects | 23,953 | 9,054,589 |
| EVA-PR: Atopic asthma + controls | 9,108 | 2,131,500 |
| EVA-PR: Atopic Asthma Only | 1,485 | 101,871 |
| EVA-PR: Controls | 289 | 1,229 |
| EVA-PR: White Blood Cell | 10,385 | 146,659 |
| EvoImmunoPop | 811 | 0 |
| Framingham Heart Study | 10,000 | 0 |
| FUSION | 0 | 36,516 |
| GTE<sub>x</sub>: BreastMammaryTissue | 6,318 | 0 |
| GTE<sub>x</sub>: ColonTransverse | 6,650 | 0 |
| GTE<sub>x</sub>: Lung | 8,686 | 0 |
| GTE<sub>x</sub>: MuscleSkeletal | 4,319 | 0 |

| **Cohort** | **cis** | **trans** |
| --- | --- | --- |
| GTE<sub>x</sub>: Ovary | 9,251 | 0 |
| GTE<sub>x</sub>: Prostate | 6,203 | 0 |
| GTE<sub>x</sub>: Testis | 2,439 | 0 |
| GTE<sub>x</sub>: WholeBlood | 3,916 | 0 |
| HELIX: Whole Blood Autosomal | 32,677 | 0 |

**Supplementary Table S1.** Sequencing Technology Implemented Across Cohorts.

| **Study/Cohort** | **Data Type** | **Technology Used** |
| --- | --- | --- |
| **EVA-PR** | RNA-seq | Illumina NextSeq 500 platform |
| **FUSION** | RNA-seq | Illumina HiSeq 2000 platform |
| **Framingham Heart Study** | RNA-seq | Illumina NovaSeq platform |
| **Colonomics** | RNA-seq | AB SOLiD System |
| **EvoImmunoPop** | RNA-seq | Illumina HiSeq 2000 platform |
| **HELIX** | RNA (extracted) | Tempus tubes; MagMAX kit; QC: 2100 Bioanalyzer and NanoDrop 1000; Infinium HumanMethylation450K BeadChip (Illumina) |
| **GTE<sub>x</sub>** | RNA-seq | Illumina TruSeq (non-stranded, polyA+ selection) platform |

**Figure S1.**
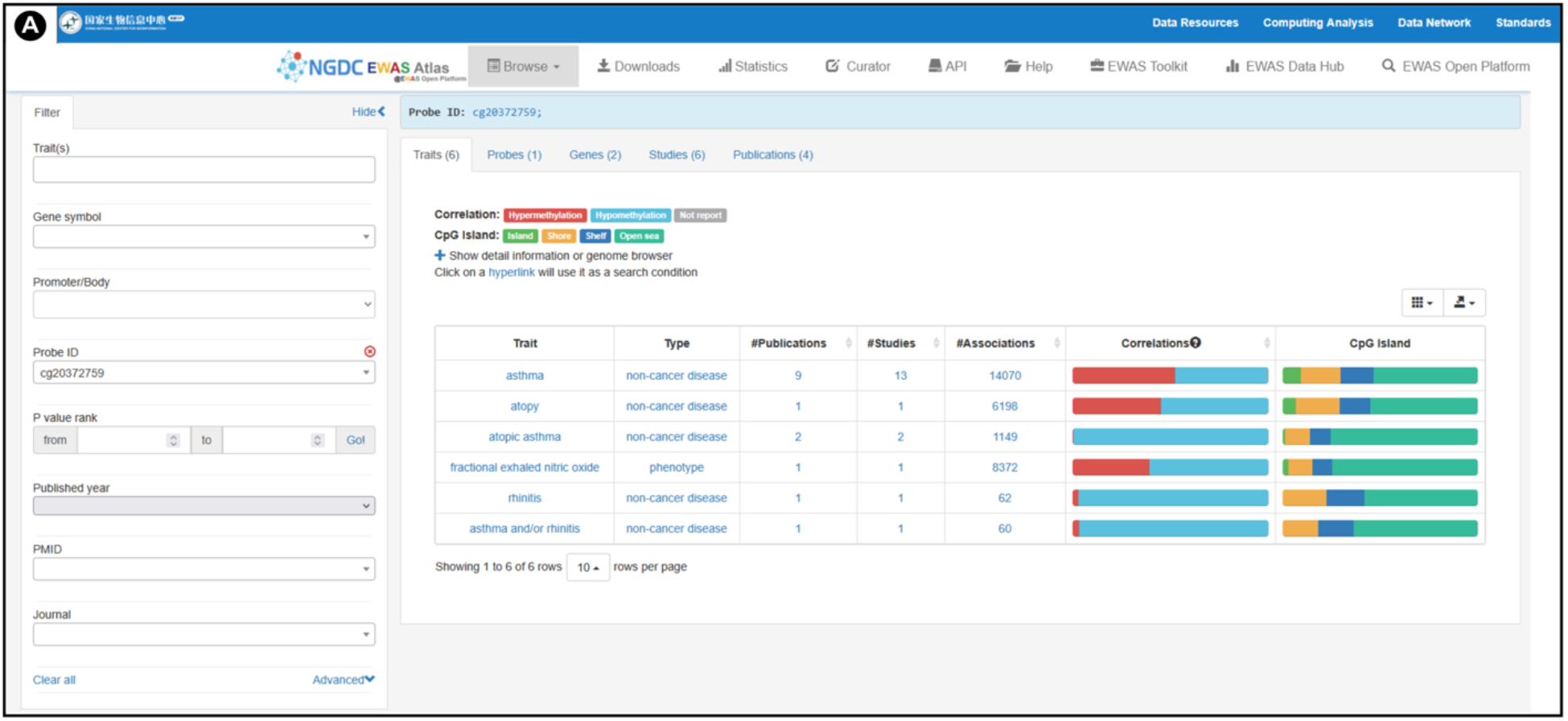
EWAS atlas results for cg20372759. Representative view from the EWAS Open Platform. Data is shown following selection of the “Open in EWAS atlas” box within the EWAS-eQTM Integration tab of Figure 3D.

## Notes

https://github.com/ads303/eQTM-Atlas

### Competing Interest Statement

The authors have declared no competing interest.

